# NAViFluX: a visualization-centric platform for interactive analysis, refinement and design of genome-scale metabolic networks

**DOI:** 10.64898/2026.01.03.697453

**Authors:** Manjunatha Beduru Krishnamurthy, Harish P S, Abhishek Subramanian

## Abstract

Genome-scale metabolic network (GSMN) models are rigorously curated, cellular-level representations of metabolism that enable flux-based metabolite fate discovery, metabolic engineering, drug target identification and context-specific multi-omics integration. However, the inherent complexity of model architectures, need for programming skills and limited visualization support restrict their broader applicability. Existing tools focus on visualization and analyses separately, necessitating tool-specific format conversions, offer either topology or flux analyses options exclusively, lack intuitive pathway-specific visualizations, database-integrated model refinement, pathway enrichment and large-scale perturbation analyses. Here, we present NAViFluX (metabolic Network Analysis and Visualization of Flux), a visualization-centric, web browser-based environment that unifies GSMN exploration through native pathway / subsystem map generation using various layouts; interactive refinement through addition of reactions from KEGG / BiGG, amending constraints, pathway merging; and modules for flux computations, topology, functional enrichment. Instead of treating visualization as a post hoc step, NAViFluX performs all actions within network views, enabling users to make decisions directly from visually interpretable subnetworks. Using three *Escherichia coli* case studies, NAViFluX is shown to characterize nutrient-specific metabolic adaptations, improve metabolic gene essentiality predictions, provide mechanistic insight into synthetic lethality and facilitate the design of an optimized carbon-fixing metabolic state, thereby democratizing GSMNs for biologically meaningful applications.

**GRAPHICAL ABSTRACT:** 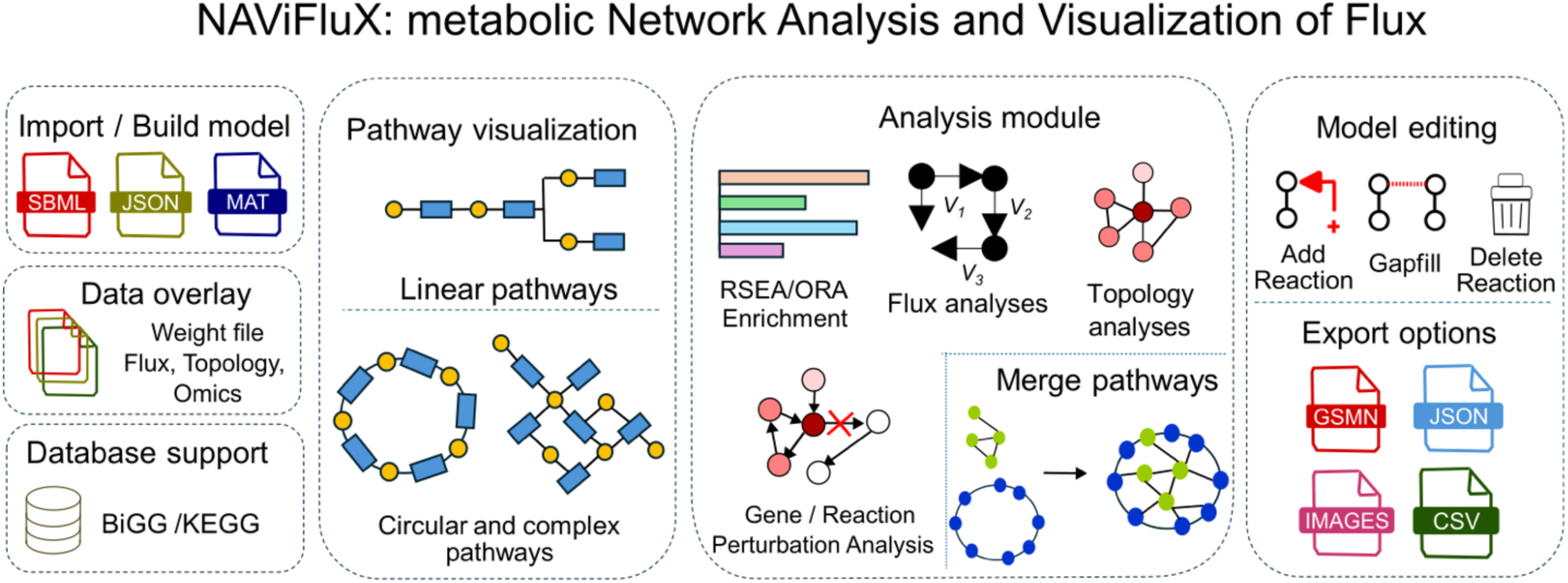

## INTRODUCTION

Genome-scale metabolic network models (GSMNs) of organisms are comprehensive metabolic reaction network libraries of gene-associated and non-gene-associated metabolic reactions that occur within any given organism [1]. GSMNs are constructed by identification and annotation of metabolic enzymes encoded within an organism’s genome and their associated reactions. Most GSMNs are rigorously curated to ensure mass & stoichiometry balance of reactions, metabolic gene / enzyme annotations, functional characterizations, metabolic pathway annotations and cross references across databases; a level of curation which even popular metabolic pathway resources like KEGG [2], MetaCyc [3] often lack. Beyond serving as static reaction repositories, GSMNs can be used for simulating various tasks like flux tracing of metabolites through metabolic pathways, optimization of cellular metabolism for bioproduction, identification of potential drug targets through in silico gene knockout simulations, and integration of multi-omics data to generate condition-, cell-, or phenotype-specific models [4].

Although GSMNs have promising applications, their utility by non-expert biologists is often limited due to the sheer complexity of the models and their organization [5]. Moreover, software like COBRA toolbox, COBRApy, COBREXA, RAVEN toolbox and CellNetAnalyzer [6], [7], [8], [9], [10] which facilitate the reconstruction and analysis of GSMNs rely on programming languages namely, MATLAB, Python and Julia. As a result, users must undergo specialized computational training for using GSMNs meaningfully. To make GSMNs more accessible, there are two groups of tools / software that facilitate metabolic network visualization and analyses - (a) GSMN - specific tools for flux analyses and / or visualization; (b) generic network graph - based tools for topology analyses & visualization.

To reduce the need for scripting, various GSMN-specific analyses and visualization tools are available for exploration. Fluxer and CAVE provide a flux balance analysis (FBA)-centered exploration of GSMNs [11], [12]. Fluxer provides flux-oriented network visualizations for tracing flow from a given metabolite towards objective functions like biomass while CAVE focuses on model editing with refinement of constraints, addition of reactions to the given GSMN and global network visualization with flow options. ModelExplorer performs structural consistency checks and manual curation of GSMNs, albeit with complex global visualizations and lack of analysis options [13]. Grohar provides automatic visualization of subnetworks upon extraction from the GSMN and predicts the effect of a knockout in the product metabolite’s neighborhood [14]. None of these tools perform topology, functional enrichment or comprehensive reaction perturbation analysis. Another tool, MetExplore uses both pre-existing metabolic networks and user-specified GSMNs for exploration, refinement, curation, omics-tailored network visualizations with flux and reaction perturbation analyses [15]. However, model analyses and network visualization are present as separate modules, treating network visualization as a post hoc step. Apart from GSMN-specific analysis tools, focused visualization tools allow the use of existing models / pathway maps from the BiGG or the KEGG database to create pathway maps, thereby bypassing the visualization limitations of these tools. Escher facilitates manual construction and editing of pathway maps using pre-drawn, template layouts on which flux or omics data can be overlaid [16]. IMFler loads Escher maps along with a GSMN to run and display flux analysis results [17]. SAMMI provides an extensive visualization package of metabolic networks supporting pathway / subsystem-specific layout visualization with omics overlays, albeit with limited network layout options and no model editing options [18]. Altogether, most GSMN-specific analysis and exploration tools put emphasis on various steps of the GSMN workflow but do not provide a comprehensive, integrated, visualization-centered workbench for performing exploration, refinement using integrated reaction databases, pathway / subsystem map generation, layout control and reaction / metabolite data overlays. Focused visualization tools, on the other hand, explicitly generate visualizations without specialized flux or topological analyses. This separation of analyses and visualization forces the generation of various output formats from the canonical GSMN SBML format to achieve cross-platform functionalities. Additionally, the existing GSMN-focused analyses and visualization tools fail to exploit the entire, curated gene, enzyme and pathway knowledgebase of GSMNs for performing complementary analyses like pathway enrichment analysis that can find differentially regulated pathways, large-scale gene-level / reaction-level perturbations and their mechanistic visualizations on metabolic networks.

Graph-based tools like Cytoscape, VisANT help in displaying unipartite or bipartite networks in various network layouts and can perform graph topology analysis to prioritize molecules with respect to their position in the network [19], [20]. Some of these tools allow integration with various omics data as well [21]. However, the user must convert GSMN models to graph network formats or edge lists which are the preferred formats for these tools. Traditionally, metabolic pathway diagrams exclude links between currency metabolites such as vitamins, cofactors, ions, water, etc., prioritizing a clean visualization to highlight biochemically meaningful transformations. None of these canonical network visualization tools are designed to support this traditional mode of visualization. Additionally, because GSMNs are genome-scale networks containing a vast number of reactions and metabolites, visualization through these tools can be often overwhelming and cumbersome. This underscores the need for pathway-level visualization which is not offered by these generic graph-based tools. These GSMN-agnostic visualization tools also do not carry the flux analysis functionalities like the GSMN-specific tools.

Given the limited availability of interactive tools that can manipulate existing reconstructions and perform diverse analyses, creation of new metabolic networks using existing reaction templates or databases becomes even more challenging. All these challenges and limitations emphasize the requirement of a robust visualization tool empowered with the ability to modify GSMNs, enable visualization-centric reconstruction and perform flux and other complementary analyses.

Here, we present NAViFluX (metabolic Network Analysis and Visualization of Flux), a visualization-centric tool with a unified environment for intuitive exploration, refinement creation and analyses of GSMNs. NAViFluX facilitates - i) native generation of metabolic pathway maps using the GSMN subsystem curations, with the choice for many intuitive linear and circular layouts, ii) specification of constraints, subsystem names, adding reactions from databases such as BiGG or KEGG, manual gap-filling and deletion of reactions on-the-fly, iii) grouping of pathway subsystems to generate broader pathways, iv) knockout simulations, topological and pathway enrichment analyses, v) mapping fluxes alongwith reaction flows, network topology metrics or relative changes of molecular abundance from any omics source onto the model-curated metabolic pathway visualizations, vi) enables the development of metabolic reaction networks from scratch using BiGG / KEGG reaction databases offering hybrid layouts that can merge subsystem-specific linear and circular visualizations and vii) the flexibility of sharing pathway maps and model files in various export formats for further manipulation in popular network / flux tools. While most tools treat network visualization as a post-hoc step, NAViFluX positions intuitive network visualization at the center of the workflow, with all model refinement and analyses revolving around / within it, allowing users to make critical decisions directly from what they see on the network.

NAViFluX was evaluated through three practical case studies on E. coli using a publicly available GSMN [22], [23]. In the first study, NAViFluX was applied to identify putative pathways responsible for the synthesis of experimentally observed metabolites across multiple carbon sources, enabling direct comparison of nutrient-specific catabolic routes through interactive pathway visualizations. In the second study, NAViFluX supported genome-scale metabolic essentiality analysis and facilitated pathway-level explanations of synthetic lethality using merged pathway views, to explain the functioning of multiple pathways in unison. In the third case study, NAViFluX was used for a metabolic engineering task in which the *E. coli* metabolic network was redesigned for optimal carbon fixation and the role of alternative carbon sources in maximizing carbon dioxide utilization towards biomass was explored.

We anticipate that NAViFluX will democratize the use of GSMNs for a wide variety of scenarios, allowing experimental and computational biologists to adopt them as a routine tool for metabolic analysis.

## MATERIAL AND METHODS

NAViFluX natively integrates visualization of GSMNs with model refinement and analyses on-the-fly, while supporting an array of input and output file formats. NAViFluX has two broad modules - “Pathway Visualizer” and “Model Builder”. The novel features of NAViFluX within these two modules are highlighted in Fig. 1. A detailed documentation entailing the features of NAViFluX, a step-by-step, hand-held guidance to use the software, tutorials for the case studies and the installation instructions are available within NAViFluX and this GitHub link https://github.com/bnsb-lab-iith/NAViFluX.

**Figure 1.**
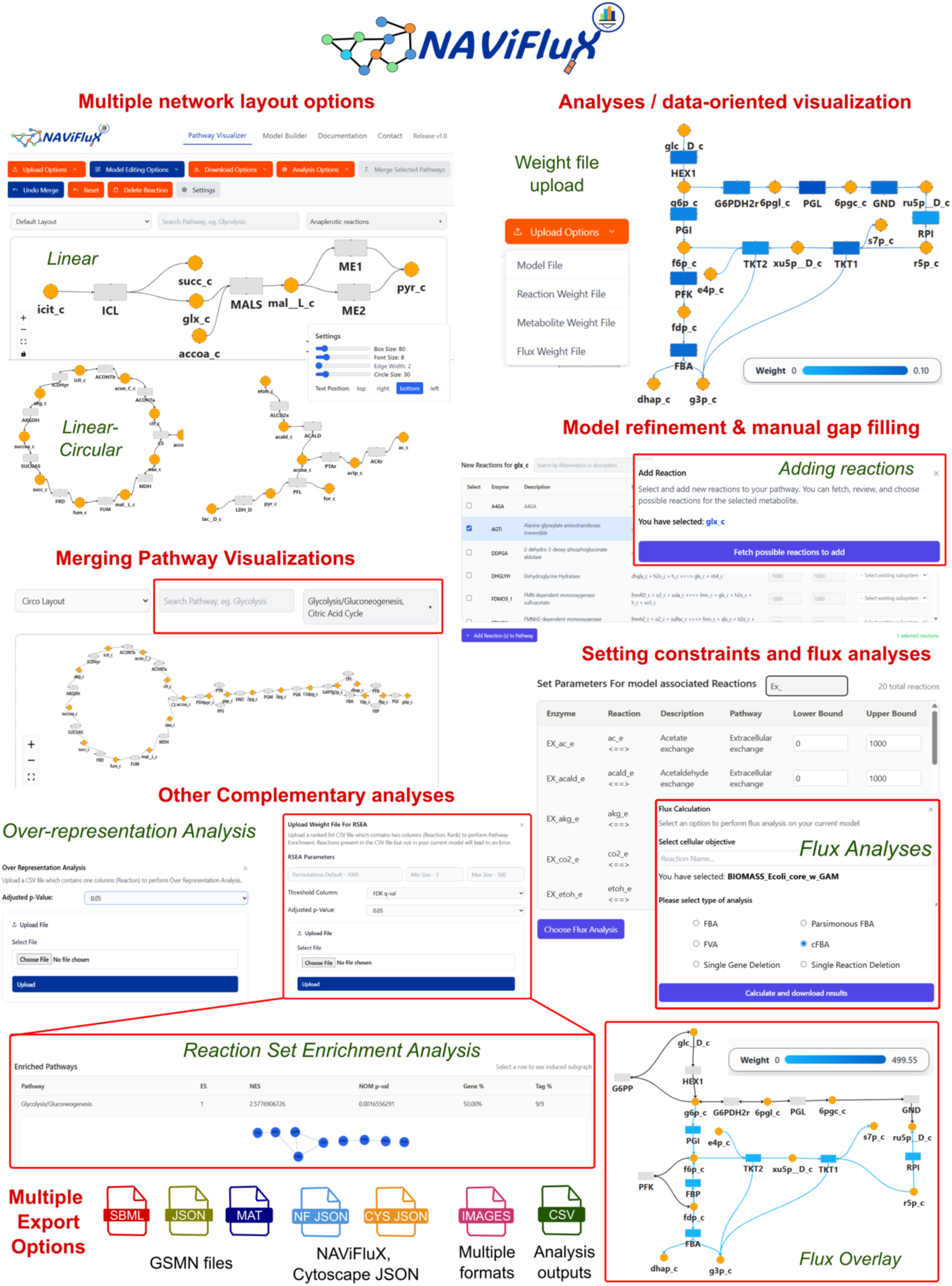
Overview of NAViFluX and its core functionalities. Schematic representation highlighting NAViFluX’s integrated workbench framework for Genome-scale metabolic network (GSMN) visualization, model refinement and analysis.

### Pathway Visualizer Module

The “Pathway Visualizer” module is the central module of NAViFluX that offers GSMN exploration, visualization, model refinement and analyses functionalities for any uploaded, BiGG- or KEGG-compliant GSMN file.

#### Uploading GSMNs and pre-processing

NAViFluX uploads GSMNs through “Upload Options”. The platform supports GSMNs in SBML, MAT or JSON formats. Currently, NAViFluX supports GSMNs generated using the KEGG or BiGG database. Once the GSMN model is loaded onto NAViFluX, the parent database (KEGG/ BiGG) is automatically detected, and the model is partitioned into pathways using the “Subsystems” field in the GSMN. A reaction-metabolite bipartite network representation is generated using the stoichiometric matrix of the uploaded model.

#### Network generation & visualization

NAViFluX generates bipartite network representations for pathways where metabolites are represented as “circular” nodes and reactions represented as “rectangular” nodes. Reaction directionality is inferred from the lower and upper bounds of each reaction specified in the model and are shown as bidirectional (reversible) or unidirectional (irreversible) edges within the network representation. NAViFluX also ensures that no currency metabolites are displayed to avoid clutter and aid appropriate visualization. NAViFluX loads the pathway-specific subnetwork onto a canvas. Nodes and edges move freely within this canvas and can be manually repositioned. The canvas also offers the options to zoom-in or zoom-out freely or to freeze the network coordinates in each position.

Edges can potentially attach to four possible positions of every node (top, bottom, left and right). To facilitate orientation of the edges between a pair of nodes, NAViFluX provides options for manipulating edge handles that can reposition edges to any of the four positions. Sizes for the reaction nodes and metabolite nodes can be distinctly adjusted for box / circle size, font size, edge width and text placement relative to node orientation providing flexibility and node-type specificity. Moreover, hovering over the reaction nodes displays information regarding the cross reference KEGG, BiGG IDs and EC number of that reaction and the weight associated with that reaction (see below). Similarly, hovering over the metabolite nodes displays information regarding ChEBI cross-reference IDs, molecular formula of that metabolite and the weight associated with that metabolite. All this information is parsed from the uploaded GSMN model file.

#### Network layout options

NAViFluX offers six different network layout options for the users to choose. The layouts can be grouped into two groups - (i) layouts enabling linear pathway visualizations: Hierarchical layouts offer linear visualizations of networks with left-to-right flow (Hierarchical-LR) and top-to-bottom (Hierarchical-BT) flow, (ii) layouts enabling hybrid linear-circular visualizations: Circo optimizes circular layouts for cyclical pathways like TCA cycle (Fig. 1). Stress layout preserves shortest path distances suitable for visualizing merged or larger pathway subnetworks. Neato is a force-directed layout that spreads nodes creating a well-spaced representation. Twopi is a radial layout that distributes nodes in concentric circles oriented using a root node (a central metabolite / enzyme).

#### Analyses / data-oriented visualizations using weight files

NAViFluX has a unique feature of overlaying heterogeneous data measurements including flux measures, network topology metrics and abundance data on reaction or metabolite nodes for obtaining pathway-level insights (Fig. 1). It is possible to upload a weight file that consists of the reaction / metabolite name and their corresponding weights (e.g. average log fold-change in gene expression of metabolic genes associated with a reaction).

Considering that every experiment measure one variable of the network in each instance (reaction, metabolite or flux), NAViFluX accepts separate weight files for reaction, metabolite weight files and flux. Uploaded weights are further standardized to make reactions / metabolites relatively comparable. Reaction / metabolite weights are mapped as color gradients to reaction / metabolite nodes. These weight files can be customized based on any source of data (e.g. omics log-fold changes, ranks, average gene expression, normalized protein and metabolite abundance, centrality, etc.), Uploading the flux weight file modifies the node colors in a gradient, based on the flux magnitude provided. Simultaneously, flux magnitudes are also mapped as colored gradients on the edges along with a change in the reversibility of arrows based on flux directions (signs) to show flux flow through the network.

#### Model Refinement

NAViFluX enables comprehensive model editing options for improving existing GSMNs. It automatically identifies whether the uploaded GSMN is a KEGG- or BiGG-compliant model file and enables model refinement using these respective databases which are natively integrated in the background. This offers useful functionalities like reaction addition, gap-filling and reaction deletion (Fig. 1). Upon selecting a metabolite, NAViFluX fetches all possible reactions related to that metabolite from the integrated KEGG or BiGG database to add reactions on-the-fly. If the user has an idea of which reaction pertaining to the selected metabolite they would like to add, the abbreviation, name, description of the reaction or metabolite can be searched from the repository. Internally, NAViFluX ensures that the reaction already present in the model is not added again, avoiding redundancy. Once the reaction that needs to be added is selected, options to set the lower and upper bounds and the subsystem label get activated. Extending this reaction addition functionality, local missing reactions can be filled by selecting any two metabolites within the canvas and adding a potential reaction from the database in which the two metabolites are involved. Any reaction within the canvas can also be deleted flexibly by applying deletion options.

#### Merging pathway visualizations

Pathway or subsystem nomenclature for reactions vary across GSMN and user preferences. Sometimes, a specific pathway name is preferred over a broader name and vice versa (e.g. Glycolysis vs. carbohydrate metabolism). This disagreement between pathway names might warrant the modification of a custom subsystem name or assignment of a new name. As a unique solution, NAViFluX offers an option to group multiple smaller subsystems into a larger subsystem using a simple selection and merging feature with the provision of assigning a new name to the merging subsystem (Fig. 1). The pathway selection feature only merges visualizations while activating the “Merge subsystems” option performs active merging within the GSMN. Additionally, all pathways in the GSMN can also be merged by selecting all pathways to get a genome-scale integrated view. With NAViFluX, users can also opt to revert to the original state if the merging procedure is not found suitable.

#### Flux Analyses

A suite of flux analyses is implemented in NAViFluX, including Flux Balance Analysis (FBA), parsimonious FBA (pFBA), Flux Variability Analysis (FVA), cycle-free FBA (cFBA), single-gene and single-reaction deletion studies. In common, all these flux analysis methodologies require the specification of flux constraints (lower and upper bounds) for the internal and exchange reactions and the selection of a cellular objective for which maximization needs to be performed (Fig. 1). Upon selection of the analysis approach in the “Flux Analyses” module, NAViFluX automatically asks for modification of reaction flux constraints. For efficiency, keyword-based search is also offered to select the reaction (by reaction, metabolite abbreviation and description) for which bounds must be updated. A flux analysis automatically overlays flux values (in case of FBA, pFBA and cFBA) and percentage reduction in growth rates (in case of single gene deletion or single reaction deletion) on the network as specified by the rules above. Simultaneously, the results are downloaded into a flux output file. This file can also be uploaded onto NAViFluX as a NAViFluX Flux weight file, whenever necessary. The flux weight file upload generates flux-based color gradients on the edges, as specified above. The Flux Analyses module generates outputs that can be imported into other programming languages for graphical and other statistical analyses.

#### Network Topology Analyses

NAViFluX also calculates standard graph-based topology metrics like degree, betweenness, closeness, eigenvector and PageRank centralities on the bipartite reaction-metabolite graph. The topology metric files are also automatically stored for further use and can also be uploaded onto NAViFluX as a reaction weight file. The weight file upload generates color gradients on the nodes based on centrality, as specified above. The “Model Network Analyses” module generates outputs that can be imported into Cytoscape and other programming languages for further graphical analyses and visualizations.

#### Functional Enrichment Analyses of reactions

There are two Functional Enrichment Analyses included in NAViFluX - (i) Reaction Set Enrichment Analysis (RSEA) and (ii) Over-representation Analysis (ORA) (Fig. 1). The subsystems curated within an input GSMN file are used for enrichment analysis thus, empowering user-oriented subsystem annotation. RSEA performs a ranking-based pathway enrichment analysis like GSEA, where reactions and their involvement in multiple subsystems is considered, instead of genes and processes. The RSEA module requires an input weight file containing reaction abbreviations and their ranks and parameters like number of permutations, minimum size (size of the subsystem with the least number of reactions) and maximum size (size of the subsystem with the largest number of reactions), the kind of multiple hypothesis testing procedure to be used (FDR, FWER) and the cut-off adjusted P-value. The RSEA output is displayed as an interactive table with Normalized Enrichment Score, ReactionRatio, Adjusted P-value and reactions enriched within each subsystem along with other essential statistics. To visualize the enriched pathway subnetworks, an enriched subsystem can be selected from the RSEA results and displayed onto a canvas within this interactive table. For performing ORA, an input list of reactions for which the subsystem enrichment analysis needs to be performed, is specified along with the adjusted P-value cut-off. Both RSEA and ORA generate output CSV files that can be imported into other programming languages for further statistical visualizations.

### Model Builder Module

NAViFlux provides a separate “Model Builder” module that enables metabolic network reconstruction from scratch using the BiGG/KEGG reaction database. Once the choice of database is selected, the interface provides two separate canvases for creating parallel pathway visualizations. Each canvas separately offers options for various network layouts. Upon creating the two subnetworks, one in each canvas, a merge feature enables merging subsystems with different layouts. While merging, NAViFluX retains the visualization layouts for each subnetwork. For instance, one can generate glycolysis, a linear pathway in one canvas using linear pathway layouts (e.g. hierarchical L-R) and TCA cycle in another canvas using cyclical pathway layouts (e.g. Circo), merge them together and make a hybrid visualization for energy metabolism retaining the individual linear and non-linear pathway visualizations. The models built in this module can be imported into the Pathway Visualizer Module for performing subsequent analyses.

#### Outputs and sharing across tools

NAViFluX offers a wide range of export formats (Fig. 1). The native output format is a NAViFluX-compliant JSON file which contains node IDs, position coordinates and model metadata. This re-usable format primarily saves the visualization state along with the fields specified in a GSMN which can be used for analyses. Certainly, common GSMN formats like MAT, SBML and COBRA-compliant JSON can also be exported for further exploration using reconstruction toolboxes like COBRApy, COBRA toolbox, COBREXA and RAVEN [6], [7], [8], [9]. The pathway / subsystem visualizations active in the canvas can be saved in multiple image formats (.png, .tiff, .svg and .pdf). Additionally, the pathway networks can be exported as edge lists of reaction pairs, metabolite pairs and reaction-metabolite pairs for import into popular tools like Cytoscape, igraph, NetworkX for specialized network analyses [19], [24]. NAViFluX also supports the export of a Cytoscape-compliant JSON file. This file retains the visualization coordinates and node / edge orientation thereby reutilizing NAViFluX visualization arrangements within Cytoscape. All analyses within NAViFluX generate outputs in CSV format.

### Technical Implementation

NAViFluX is implemented as a web-based client-server system with a React-based frontend that runs on a browser and a Flask framework-based backend server that handles analyses modules and functions. The frontend developed using React v.18.2 with the Vite build system hosts the graphical user interface, while the backend developed using Flask v.3.1.2 executes the various computational analyses. ReactFlow v.11.11.4 was used for interactive graph visualization and generation of NAViFluX- and Cytoscape-compliant .json files. The backend scripts for implementing analyses were written in Python v.3.13. COBRApy v.0.30 package was used for reading and parsing GSMN files, conversion of model file formats and performing constraint-based analyses [6]. GLPK solver was used for performing flux analyses. Communication between the frontend and backend was achieved through RESTful APIs, with Flask-CORS v 6.0.2. enabling cross-origin resource sharing. Network layout generation is supported by the Dagre javascript library v0.8.5 for hierarchical layouts and ELK.js library v0.10 for layered graph visualization. Network topology analyses and edge list generation were performed using NetworkX v 3.6.1. Functional enrichment analyses like RSEA and ORA were customized for reactions using GSEApy v1.1.11 [25]. All the scripts underlying NAViFluX build, and implementation were written in-house.

### Additional analyses for case studies

All the core analyses like flux analyses, gene / reaction perturbations, ORA and flux-oriented visualizations were performed within the NAViFluX environment. For case studies 1 and 2, growth media compositions reported in corresponding studies (M9 and MOPS medium + corresponding carbon source) was used to infer the constraints of nutrient uptake using MetaboTools functions implemented in COBRA toolbox v.3.0 [7]. All data preprocessing and analyses were performed using the *tidyverse*, with *dplyr* for data manipulation. Unless stated otherwise, visualizations were generated using *ggplot2*, while heatmaps in the case studies were produced using the *ComplexHeatmap* package [26]. All these packages were implemented in R v.4.4.3.

## RESULTS

### Comprehensive benchmarking of NAViFluX against available GSMN modification, analyses and visualization tools

Several GSMN-specific tools are publicly available in assisting metabolic network visualization, analyses and refinement, albeit, with significant limitations in scope and flexibility of use. Broadly, there are four sets of tools that can work on GSMNs - (i) exclusive analyses-oriented tools which perform analyses on GSMNs (e.g. FLUXestimator [27]); (ii) exclusive visualization-oriented tools which generate network layouts and overlay flux or omics data on networks (e.g. Escher, IMFLer, SAMMI, CytoSeed, FLUXer, Grohar); [12], [14], [16], [17], [18], [28] (iii) exclusive model manipulation tools (e.g. ModelExplorer [13]); (iv) hybrid tools which perform model manipulation, analyses as well as visualization (e. g. CAVE, MetExplore V2) [11], [15]. In comparison, NAViFluX provides an integrated workbench that offers novel features for model manipulation, analyses and visualization beyond what is implemented in previous tools. Comparison of NAViFluX features with other tools are provided in Fig. 2.

**Figure 2.**
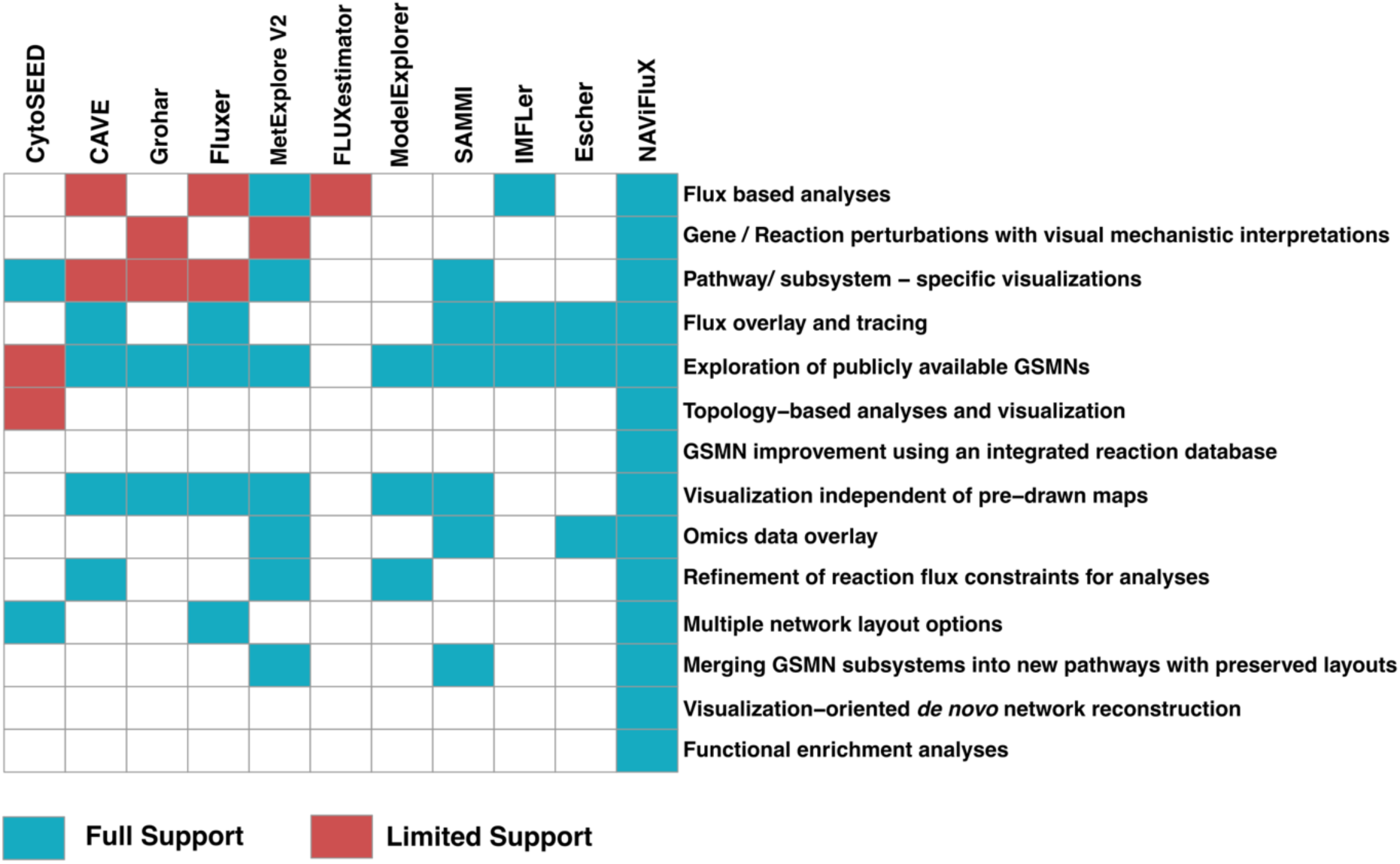
Comparison of NAViFluX features with other GSMN development, analysis and visualization tools. Tools providing full support for certain actions are colored in blue whereas tools providing partial support for those functions are colored in red.

#### GSMN manipulation and modification

Except FLUXestimator, all other tools support exploration of publicly available GSMNs. Some tools provide limited support like CytoSEED which offers visual exploration of only ModelSEED GSMNs. Reaction flux constraints can be refined using exclusive model manipulation or hybrid tools like ModelExplorer, CAVE and MetExplore V2. Also, customized reactions can be defined in these tools. However, all these tools offer network visualization only as a post hoc step for display of reactions as a subnetwork or flux overlays. Also, none of these tools offer database-integrated model modifications. None of the tools except SAMMI and MetExplore V2 offer merging of pathway subsystem visualizations.

However, SAMMI and MetExplore V2 do not make active modifications in the model while merging the subsystems, rather use it solely for merging visualizations. In contrast to all these tools, NAViFluX offers an interactive, visualization-centric approach which allows better visual control of the GSMN helping with decision-making at every step. Additionally, NAViFluX is integrated with the BiGG and KEGG reaction database thereby facilitating addition of database reactions to the uploaded GSMN instead of manually defining the reactions. NAViFluX also offers manual refinement of constraints for any reaction in the GSMN using an interactive table that is integrated with text-based search. Instead of scrolling through the list of reactions, reactions can be filtered based on reaction names, metabolite names and abbreviations. Moreover, the upper and lower flux bounds can be iteratively set. Another interesting feature of NAViFluX is that it can merge pathway subsystems to generate new pathways along with a delete reaction option thereby providing flexibility in creating new pathways. While merging subsystems, NAViFluX actively modifies the GSMN. These features make NAViFluX uniquely useful for model manipulation. Using the Model Builder module, NAViFluX also facilitates *de novo* visualization-oriented network construction which no other tool can perform.

#### Analyses options

Except for visualization-oriented tools like SAMMI, Escher, CytoSEED and ModelExplorer, all other tools natively provide only a few flux analysis options. FLUXestimator exclusively performs single-cell or bulk transcriptomics - oriented flux estimation on GSMNs (scFEA). In contrast, NAViFluX offers options for a wide variety of flux analyses like FBA, FVA, pFBA and cFBA. Similarly, none of these tools natively perform network topology analyses to find bottlenecks or central reactions in metabolic networks. CytoSEED, being a Cytoscape plugin for ModelSEED GSMNs, can be tuned so that network analyses can be performed within Cytoscape. However, it is important to note that the CytoSEED plugin is now defunct. NAViFluX on the other hand, can perform network centrality analyses to estimate metrics like degree, betweenness, closeness, PageRank and eigenvector centrality to facilitate interpretation of network organization. None of these tools perform gene / reaction level perturbations comprehensively. For instance, Grohar, MetExplore V2 perform perturbations of a single reaction and observe its effect on local reactions or a selected objective function. However, a comprehensive knockout (KO) analysis that can also measure the relative effect of the KO on a defined objective function is only available in NAViFluX. As NAViFluX offers options to overlay any weight file and interactive network refinement, the KO analysis can be coupled with flux visualizations for mechanistic interpretations, as demonstrated in Case Study 2. Functional enrichment analyses like ORA or RSEA that utilize the GSMN subsystem information are uniquely implemented in NAViFluX, thereby valuing model pathway curation.

#### Visualization options

All the specified tools except FLUXestimator provide network visualization options. CytoSEED, Escher and IMFLer rely on pre-drawn maps and thus, are limited to only GSMNs for which maps are available in KEGG or Escher JSON format. MetExplore V2, SAMMI and Escher provide omics overlay options on reactions while CAVE, Fluxer, SAMMI, IMFLer and Escher provide overlay of flux magnitude on reaction nodes and the directionality on the edges connecting them. None of the tools offer topology-based visualization on networks. SAMMI, CytoSEED and MetExplore V2, exclusively provide full-fledged pathway-specific visualization. CAVE, Grohar and Fluxer can generate subnetworks based on the specification of source and target reactions which can work only as a means for post analyses interpretations. Apart from all these features, only CytoSEED and Fluxer offer multiple network layout options. As NAViFluX prioritizes visualization, it implements all these important features including multiple layout options, overlay of omics, topology, flux and reaction perturbations on reaction nodes and merging of pathway subnetworks to generate new pathways / subsystems while retaining their original pathway layouts. In addition, NAViFluX also supports native, currency metabolite-free pathway-specific visualization, addition and deletion of reactions by interactively selecting metabolites / reactions in the network display canvas and node and edge modification options with information displays, thereby prioritizing biochemically intuitive visualization.

Overall, NAViFluX encompasses the most essential functionalities of model modification, analyses and visualization required for conducting computational systems biology experiments on metabolic systems. NAViFluX is thus comparable to visualization tools like SAMMI and CytoSEED and model analyses and manipulation tools like CAVE and MetExplore V2 while offering more intuitive control to the user.

### Case Studies

Here, we present three case studies that demonstrate the benefits of NAViFluX in performing various analyses and facilitating the interpretation of metabolic mechanisms through network visualizations. The case studies showcased here represent small example projects. It is important to note that the results demonstrated here represent NAViFluX’s functionality and may or may not converge to a full-fledged hypothesis.

#### Case study 1: Condition-specific flux analyses and visualizations for the comparison of carbon source utilization by Escherichia coli

To demonstrate NAViFluX’s capability for obtaining flux-level insights about metabolism, we exploited a targeted metabolomics dataset for uncovering carbon source-specific metabolic adaptations in *Escherichia coli* [29]. The original study measured 101 primary metabolites across 19 environmental conditions in *E. coli BW25113* to investigate the variations in relative metabolite abundances under various nutrient sources and stresses. Out of the 19 environmental conditions, we chose metabolomics profiles of 11 carbon source conditions for comparison. The *E. coli* iJO1366 model was simulated for growth on 11 carbon sources independently to predict potential utilization pathways (Fig. 3A). NAViFluX FVA showed that the optimal growth rate varied between 0.18 mmol/(gDW.hr) to 0.51 mmol/(gDW.hr) depending upon the carbon source used for simulation. The growth rate of *E. coli* under glucose was predicted to be 0.51 mmol/(gDW.hr) which is also reported in previous studies [30].

**Figure 3.**
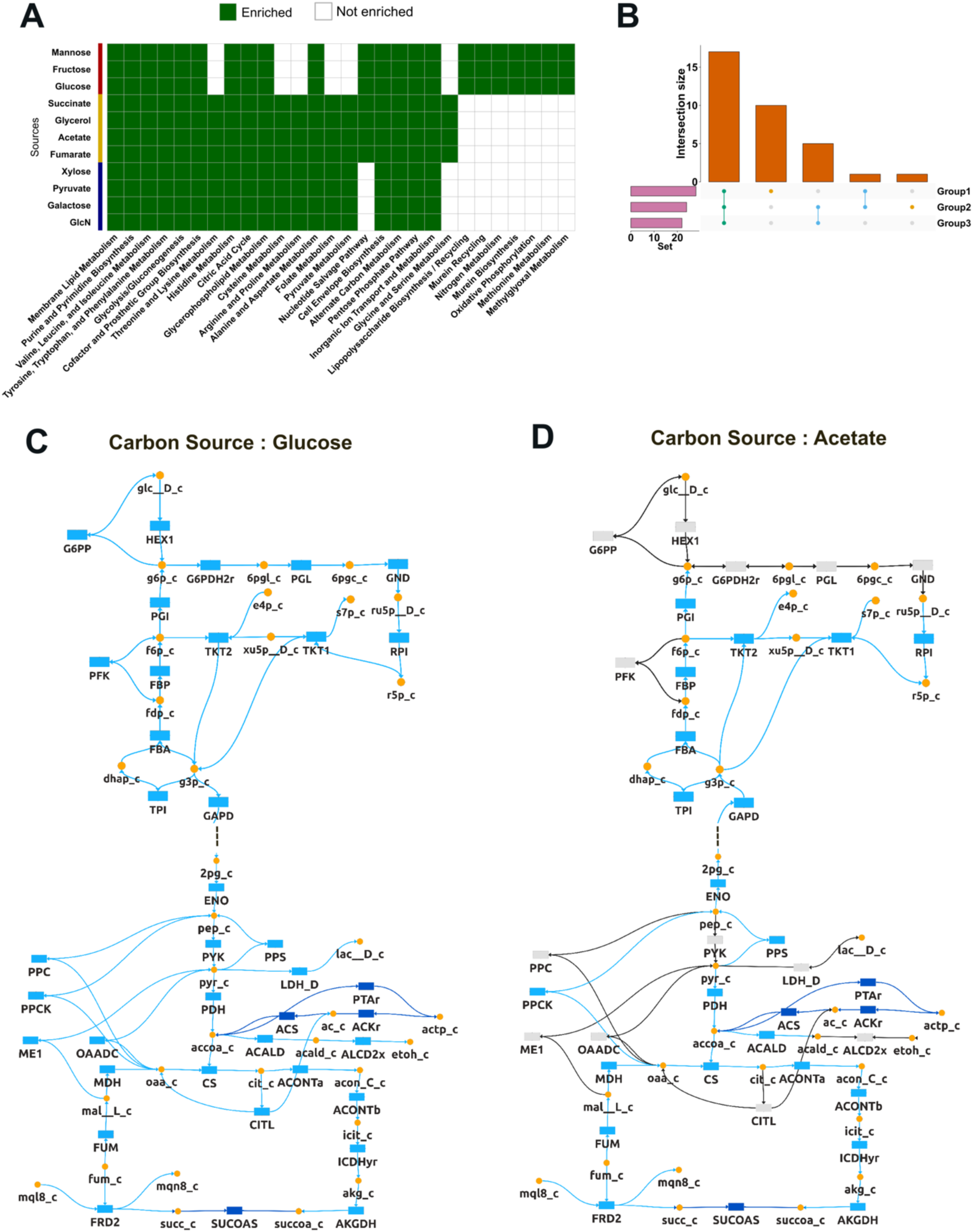
NAViFluX predicts interpretable metabolic adaptations of *Escherichia coli* to carbon sources. (A) Heatmap summarizing over-representation analysis (ORA) of flux-active pathways as obtained from flux variability analysis (FVA) across 11 carbon source conditions. Green indicates enriched pathways, while white denotes non-enriched pathways. Conditions cluster into three distinct enrichment groups (Groups 1–3), reflecting shared and unique pathway activation patterns across carbon sources. (B) UpSet plot showing the intersection of enriched pathways among the three groups, highlighting pathways common to all groups as well as those uniquely enriched in individual groups. (C) Flux visualization of a merged central carbon metabolism subnetwork for the representative carbon source - glucose (Group 1). Colour gradient corresponds to the average of minimum and maximum FVA flux values. Growth on glucose exhibits a high flux glycolytic state with active oxidative, non-oxidative pentose phosphate pathways and lactate production. (D) Flux visualization of a merged central carbon metabolism subnetwork for the representative carbon source - acetate (Group 2). Growth on acetate shows a strictly gluconeogenic state with an active TCA cycle, reliance on non-oxidative PPP and no lactate release.

The iJO1366 *E. coli* model was uploaded into NAViFluX and the estimated uptake constraints were added to the model using the “Flux Analyses” module. Setting biomass as the objective function to be maximized, FVA within this module was used to obtain “flux-active” reactions (reactions with non-zero minimum or maximum rates) (Supplementary Table S1). FVA captures the allowable flux ranges that can produce maximal biomass. As expected, metabolites tested in the experiments were a subset of these predicted flux-active metabolites (metabolites associated with the flux-active reactions) (Supplementary Figure S1, Supplementary Table S1). This suggested that the metabolites chosen in the experiment were growth-associated metabolites and the inferred fluxes within the model effectively represented each environment’s entire metabolic potential.

The active reactions in each environmental condition were then subjected to ORA using the “Functional enrichment analysis” module of NAViFluX to obtain statistically over-represented processes. Exporting the over-representation outputs for each condition, a heatmap for comparing the enriched and non-enriched processes in each condition was generated. The heatmap revealed three clusters (Groups 1, 2 and 3) of enrichment profiles across conditions suggesting that the catabolic pathways overlap between certain carbon sources (Fig. 3A). 10 unique pathways in Group 1, one unique pathway in Group 2 and 17 pathways common to all the three groups were found (Fig. 3B, Supplementary Table S2). Pathways common across clusters, as well as those unique to individual groups, were identified and these analyses demonstrated that different carbon inputs induce specific activation patterns within central metabolic, anaplerotic, and biosynthetic pathways.

With the aim of comparing flux distributions between Groups 1 and 2, a merged pathway subnetwork by merging 4 pathways was generated in NAViFluX. The flux distributions (average of the minimum and maximum FVA fluxes) of the two representative input carbon sources (glucose from Group 1 and acetate from Group 2) were overlaid onto this merged pathway subnetwork using flux weights file generated from FVA analysis in NAViFluX. Growth on glucose resulted in a high glycolytic flux state, characterized by strong activity in both upper and lower glycolysis and oxidative and non-oxidative branches of the pentose-phosphate pathway for nucleotide and amino acid biosynthesis (Fig. 3C). The high glycolytic flux increased overflow metabolism (high lactate, acetate and ethanol production) by coupling to the TCA cycle to maintain NAD-NADH redox balance. An activation of citrate lyase (CITL) reflecting cytosolic acetyl-CoA production to reroute excess glucose-carbon flux was also observed.

In contrast, growth on acetate revealed a strictly gluconeogenic state, with carbon flowing upward from the TCA cycle to produce hexose phosphates suggesting the ultimate requirement of glycolytic / PPP intermediates for biomass precursor synthesis (Fig. 3D). Flux through upper glycolysis was absent and glucose-6-phosphate was exclusively catabolized through non-oxidative PPP. Additionally, growth on acetate shows the generation of NADPH exclusively through isocitrate dehydrogenase (ICDHYr). Oxidative PPP is inactivated probably to prevent carbon loss through CO_2_ release suggesting acetate to be a poor carbon source. NAViFluX network visualization shows an active TCA with no overflow metabolism. Supporting this, auxiliary enzymes like PEP carboxylase (PPC) and citrate lyase (CITL) remained inactive as the TCA cycle was able to maintain acetyl-CoA and oxaloacetate pools at sufficient levels, obviating the need for additional carbon-fixing or splitting reactions.

Overall, this case study demonstrates NAViFluX’s ability to tune model constraints, perform flux analyses, over-representation analysis and make flux-weighted network visualizations.

#### Case study 2: Improving essential gene predictions in Escherichia coli using intuitive single-gene deletion simulations

NAViFluX can also be used for predicting essential metabolic genes and uncovering the metabolic reason (mechanism) underlying their essential nature. We evaluated NAViFluX’s ability to predict essential metabolic genes by comparing its predictions with essential genes reported in the KEIO single-gene deletion library for *E. coli* K-12 MG1655 [31]. This reference library contains information of 3,985 known nonessential genes and 303 known essential genes.

First, we compared whether there are flux differences between the reported essential and non-essential reactions (reactions corresponding to the reported genes) in the wild-type model. For this purpose, cFBA was performed using the “Flux Analyses” module in NAViFluX by constraining the metabolite environment based on the MOPS growth medium (Materials and Methods). cFBA estimates reaction fluxes that can maximize biomass by correcting for futile cycles. cFBA was performed using the “Flux Analysis” module of NAViFluX and reaction fluxes were saved. Interestingly, non-essential reactions carry larger flux magnitudes as compared to essential reactions suggesting that non-essential reactions within the *E. coli* metabolic network are largely involved in flexible, redundant routes that can be reconfigured upon knockout of a non-essential reaction (Fig 4A). Essential reactions on the other hand, sit as metabolic “chokepoints” in more linear parts of the network.

**Figure 4.**
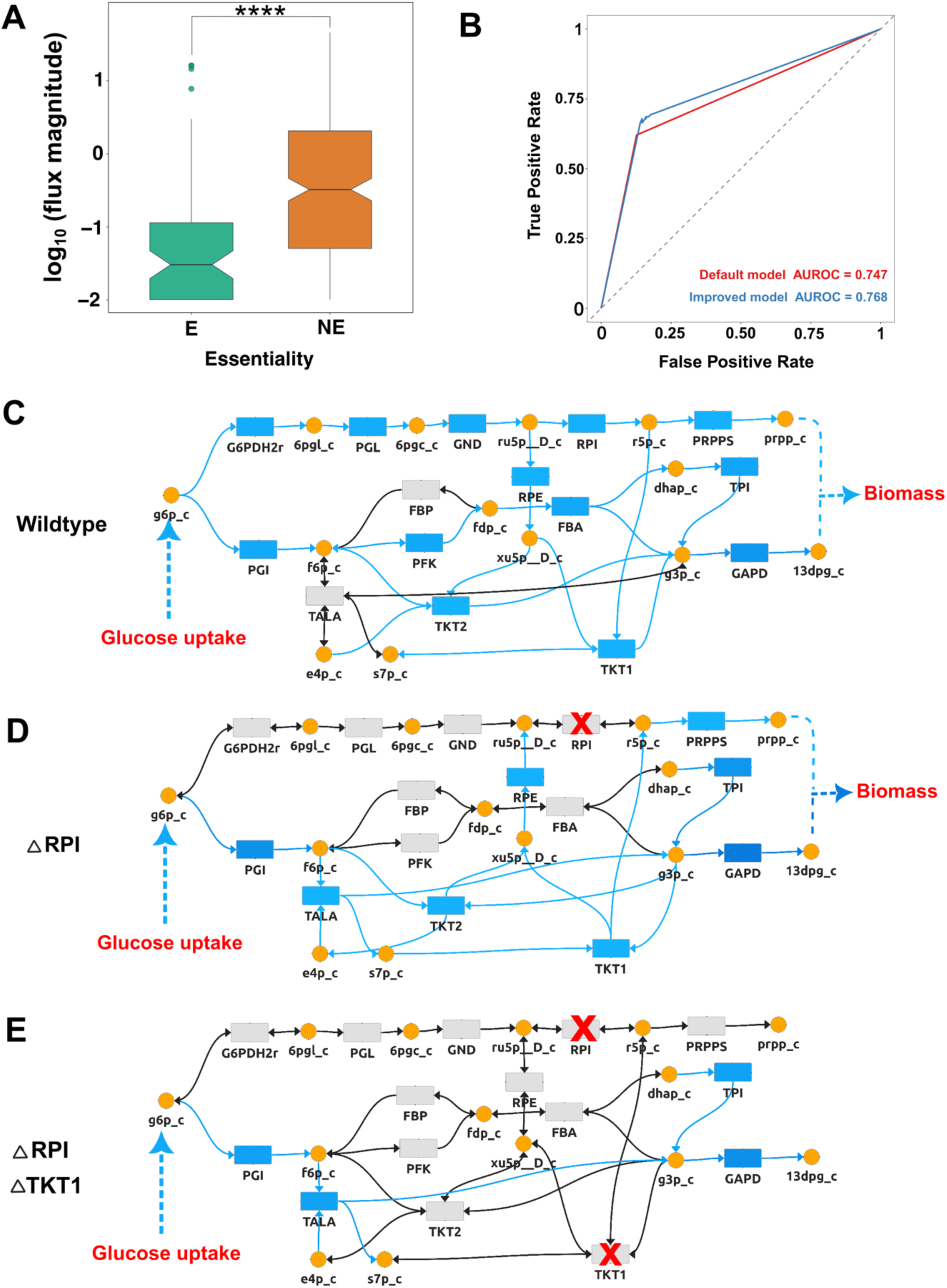
NAViFluX based prediction of metabolic gene/reaction essentiality and mechanistic interpretation through flux visualization. (A) Comparison of reaction fluxes in the wild-type *E. coli* iJO1366 GSMN between reactions associated with the experimentally essential (E) and non-essential (NE) genes as reported in the KEIO single-gene deletion library. Plotting is performed in log-scale of absolute flux. Overall, non-essential reactions exhibit significantly higher fluxes than essential reactions (Wilcoxon test, p = 4.36 × 10⁻¹¹). (B) Receiver operating characteristic (ROC) curves comparing essential gene prediction performance using default model constraints (red; auROC = 0.747) and minimally refined internal constraints (blue; auROC = 0.768), demonstrating improved prediction performance upon constraint refinement. Constraint refinement and genome-scale gene knockouts were implemented in NAViFluX. (C) Flux visualization in the *prpp* synthesis subnetwork under wild type conditions. Cycle-free flux balance analysis (cFBA) indicates that *prpp* synthesis is preferred via the oxidative pentose phosphate pathway mediated by ribose-5-phosphate isomerase (RPI). (D) Flux visualization in the *prpp* synthesis subnetwork upon single reaction knockout of RPI in the *E. coli* iJO1366 model. (E) Flux visualization in the *prpp* synthesis subnetwork upon combinatorial knockout of RPI and TKT1

The “Single Gene Deletion” functionality within the “Flux Analysis” module of NAViFluX was performed to predict essential genes. Comparison of the predicted knockout-growth rates with the KEIO essentiality yielded an area under the receiver operating characteristic (auROC) of 0.74, indicating that the iJO1366 model with default constraints had a reasonable ability to predict essential genes in *E. coli*. We hypothesized that accurate prediction of essential genes is largely dependent on internal network constraints specified within the iJO1366 GSMN. For instance, phosphoribosyl pyrophosphate synthetase (PRPPS) is an essential enzyme for the synthesis of *E. coli* biomass [31], [32]. The default constraints within the iJO1366 model predicted PRPPS to be non-essential. Exploring the model in NAViFluX revealed that the Ribose 1, 5 bisphosphokinase (R15BPK) reaction is involved in an alternative catabolic route to produce phosphoribosyl pyrophosphate (*prpp*) thus rendering PRPPS to be redundant and non-essential. R15BPK is a dispensable enzyme in *E. coli [33]*. Using NAViFluX, we constrained the R15BPK reaction flux to 0 mmol / (gDW.hr) thereby eliminating the alternative route so that PRPPS would become a preferred route for biomass. Similarly, periplasmic transport of D-Glucose (GLCptspp) and hexokinase (HEX1) were both considered as alternative sources for glucose-6-phosphate formation and diversion of glucose towards ribose-5-phosphate synthesis. To keep both GLCptspp and HEX1 reactions active in the GEM model, the upper bound of GLCptspp was constrained to 0.5 mmol / (gDW.hr) such that HEX1 compensates for the remaining requirement of catabolizing glucose for *r5p* synthesis. Applying these minimal changes to the GSMN, slightly increased the model performance (auROC=0.76) and ensured that PRPPS was essential for nucleotide and amino acid synthesis (Fig. 4B). This established NAViFluX’s ability to perform single gene deletion simulations and revision of flux constraints within the model for the improvement of single gene essentiality prediction.

It is well-known that catabolism of glucose for *prpp* synthesis can potentially be achieved through the oxidative and non-oxidative pentose phosphate pathways (oxPPP and non-oxPPP). To study this aspect, NAViFluX’s functionality of merging subsystems was exploited. Glycolysis, PPP and histidine metabolism subsystems from the iJO1366 GSMN were merged to obtain a net glucose catabolic pathway. This subnetwork was named as the *prpp* synthesis subnetwork (Fig. 4C). Reactions essential for explaining *prpp* synthesis were retained in this merged subsystem. This network clearly showed the two parallel metabolic paths for glucose catabolism into biomass synthesis, involving ribose-5-phosphate isomerase (RPI) and transketolase I (TKT1) reactions which uniquely represent catabolic checkpoints for oxidative (glucose-6-phosphate (*g6p*) to *prpp* via D-Ribulose 5-phosphate (*ru5p_D_c*)) and non-oxPPP (glucose-6-phosphate (*g6p*) to *prpp* via sedoheptulose-7-phosphate (*s7p*) and glyceraldehyde-3-phosphate (*g3p*)) respectively. This visualization was stored in NAViFluX’s native JSON format for overlaying the wild-type and knockout fluxes. Performing cFBA in the wild-type situation shows that RPI-mediated oxidative PPP is the preferred pathway for *prpp* synthesis (Fig. 4C).

We investigated whether RPI is essential or not, by fixing the upper and lower bounds of RPI to 0 mmol/ (gDW.hr) simulating the effect of a single reaction knockout within the system. Upon RPI knockout, *prpp* synthesis becomes dependent on non-oxPPP for sustaining glucose catabolism (Fig. 4D). The network visualization revealed that TKT1 mediated non-oxPPP activity. By constraining upper and lower bounds of RPI and TKT1 reaction to 0 mmol / (gDW.hr), a double knockout simulation was performed in NAViFluX. It can be observed that both oxidative and non-oxidative PPP are unable to catabolize glucose towards *prpp* synthesis, thus rendering the double knockout lethal (Fig. 4E). As expected, the combination of RPI and TKT1 was essential for biomass via *prpp* synthesis. This exercise showcases the ability of NAViFluX in visually guiding interventions and conducting metabolically meaningful perturbations.

Overall, this case study demonstrates that NAViFluX performs tunable gene essentiality predictions and can uncover visual, mechanistic insights into redundancies and vulnerabilities within metabolism.

#### Case study 3: Engineering an Escherichia coli metabolic network for carbon fixation

To illustrate NAViFluX’s utility in simulating engineered metabolism, we considered a study where *E. coli BW25113* strain was engineered to synthesize sugars solely from CO_2_ via incorporation of essential Calvin-Benson-Bassham (CBB) cycle enzymes, thereby, leading to a hemiautotrophic growth switch [34]. We attempted to reproduce the results of this study on the iJO1366 model and extend it further to identify carbon sources that can couple with the engineered CBB cycle to maximize CO_2_ fixation. Like case studies 1 and 2, the growth media used in their study (M9 medium) was used to define uptake constraints. To reproduce their study, we generated the “mutant” metabolic network by constraining the reactions associated with the genes reported to be deleted in their study to zero flux (phosphoglycerate mutase PGM, glucose-6-phosphate dehydrogenase G6PDH2r, Phosphofructokinase PFK and the glyoxylate shunt reactions Isocitrate lyase ICL, Malate Synthase MALS and Isocitrate dehydrogenase ICDHx) and the addition of CBB enzymatic reactions RuBisCO and phosphoribulokinase (RBPC and PRUK) using NAViFluX. The “Model Editing Options” module adds these reactions from the BiGG database integrated within NAViFluX. This action simulates the knockout of an organism’s genes and the expression of novel recombinant CBB enzymes which is required to ensure the activity of an engineered pathway that can fix CO_2_ towards biomass synthesis.

The model was further constrained by limiting the pyruvate uptake flux to 16 mmol/ (gDW.hr), as reported in the study [34]. cFBA was performed on these constraints using the “Flux Analyses” module of NAViFluX and a flux profile was generated. A growth rate of 23 mmol / (gDW.hr) was achieved, matching with previous experimental and GSMN studies [34]. Tracing the pathway fluxes through the GSMN, the experimentally engineered division of central carbon metabolism into two distinct functional modules could be reproduced (Fig. 5A). This division disrupts the flow of carbon, generating distinct functional modules (also called flux modules) - the disjoint CBB and Energy cycles, as observed from the flux-overlaid networks. The upper glycolysis pathway and the PPP constitute the CBB module which is exclusively involved in CO_2_ fixation. Carbon enters this module via Ribulose-1,5-bisphosphate carboxylase (RBPC) and phosphoribulokinase (PRUK). This module allows the regeneration of intermediates and synthesis of the hexose, pentose, and triose phosphates required for biomass synthesis from CO_2_. In contrast, the energy module being uncoupled from the CBB module with respect to carbon flow, utilizes pyruvate exclusively for ATP synthesis and supplies reducing equivalents for CO_2_. The separation of carbon flow between energy generation and carbon fixation establishes a partial CO_2_-dependent growth called hemiautotrophic growth.

**Figure 5.**
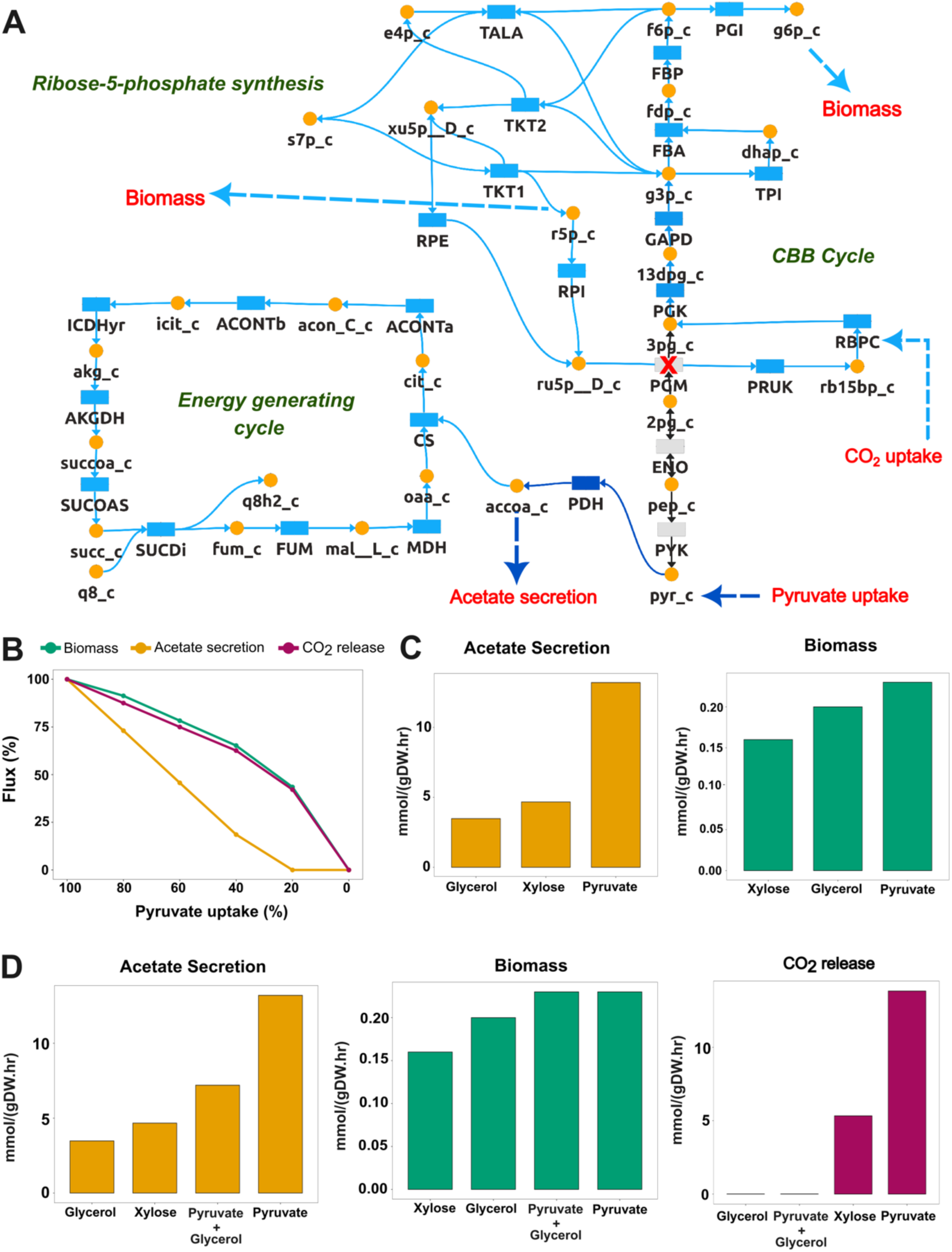
NAViFluX-enabled simulation of hemiautotrophic growth in engineered *E. coli* and its optimization. (A) Flux visualization of pyruvate fate in the engineered *E. coli* iJO1366 model simulated for hemiautotrophic growth. NAViFluX analyses and visualization reveals the emergence of two disjoint but redox-coupled functional flux modules: a CBB module dedicated to CO₂ fixation and precursor biosynthesis, and an energy generating module fueled by pyruvate oxidation. (B) Effect of progressively reducing pyruvate uptake on biomass formation, acetate secretion, and CO₂ release. (C) Comparison of acetate secretion and biomass production under individual carbon sources (glycerol, xylose, and pyruvate). (D) Maximization of CO_2_ fixation into biomass with minimal overflow metabolism.

Despite carbon flow being decoupled between the two modules, the energy and the CBB module are redox coupled. Although the above simulations with pyruvate uptake at 16 mmol/(gDW.hr) reflect the intended hemiautotrophic behavior, an excessive acetate secretion also becomes apparent, suggesting an excessive utilization of pyruvate to meet the redox requirements of the CBB module (Figs. 5A and B). Additionally, it can be observed that there is still a net CO_2_ release under optimal pyruvate uptake. An ideal optimal condition should capture maximum CO_2_ fixation and minimize CO_2_ release. Moreover, excessive acetate generation might lead to increased acidity in the environment and can be detrimental to cell growth [35]. Using NAViFluX, multiple pyruvate uptake constraints were fixed in the model, and the rates of CO_2_ and acetate release were recorded. Consistent reduction of pyruvate decreases acetate secretion and at a pyruvate uptake rate of 3.2 mmol/(gDW.hr), which is roughly 20% of its optimal rate, acetate release becomes zero. Below this pyruvate uptake threshold, acetate secretion rate becomes zero and is no more correlated with changes in pyruvate uptake (Fig 5B). With further reduction in pyruvate uptake, a successful decrease in overflow metabolism was achieved. Similarly, net CO_2_ release becomes zero suggesting a complete utilization of CO_2_. Although this is the intended result, there is an important caveat, growth rate is halved.

The “Flux Analysis” module of the NAViFluX was again used to generate simulations on individual carbon sources and their combinations, one at a time. As observed in Fig. 5C, growth in glycerol demonstrated the least acetate secretion rate as compared to xylose and pyruvate, even though pyruvate is the preferred carbon source for biomass synthesis. We hypothesized that glycerol could be an alternative carbon source that can maximize CO_2_ fixation (minimize CO_2_ release) while still maintaining optimal *E. coli* biomass synthesis. Compared to the network simulated on individual carbon sources - xylose, glycerol and pyruvate, a combination of glycerol and suboptimal pyruvate (fixing pyruvate uptake = 4, glycerol uptake = 7 mmol/gDW/hr) restores biomass equivalent to the default pyruvate uptake while halving acetate release (Fig. 5D). Interestingly, in this situation, CO_2_ release is nearly eliminated, very similar to the glycerol-only situation suggesting that this combination achieved the best of both worlds - maximal biomass and CO_2_ fixation.

With this case study, we were not only able to reproduce their pathway / functional module observations but also extend the analyses to intuitively achieve optimal CO_2_ fixation into biomass, a novel application of NAViFluX visualization and flux analyses.

## DISCUSSION

NAViFluX enables an integrated, visualization-centric environment that allows entry-level users to perform complex analyses such as flux analyses, pathway enrichment analyses, data-oriented visualizations, reaction- and pathway-level editing of publicly available / reconstructed GSMNs and intuitive network visualizations that facilitate hypothesis generation & interpretations. The main philosophy behind NAViFluX is to eliminate the traditional separation between mathematical analyses and network visualizations by allowing users to manipulate constraints, add or delete reactions, merge pathways, run various analyses while immediately inspecting results on the merged pathway subnetworks. NAViFluX thus shifts GSMN usage from script-driven workflows to visually guided, iterative reasoning about metabolic network organization, flux distributions across pathways and essential reactions. Comparing NAViFluX with existing metabolic network visualization and analyses tools highlights key features that makes NAViFluX unique. Features like native pathway / subsystem map generation directly from the GSMNs with pathway-specific network layouts, interactive reaction-level editing by directly accessing BiGG or KEGG reaction reference databases, *de novo* visualization-oriented GSMN model building, unique functional enrichment and topology analyses options and exportable models and maps that can be transferred for further modifications / analyses using other tools make NAViFluX a unique workbench for comprehensive metabolic network analyses. Case study 1 illustrates the potential of NAViFluX in visually distinguishing glycolytic versus gluconeogenic states and different uses of the pentose phosphate pathway under glucose and acetate growth by overlaying flux profiles on merged pathway subsystems. NAViFluX identifies the coupling between various pathways providing mechanistic interpretations for why certain carbon sources are preferred and how that might help achieve optimal metabolic outputs (e.g. biomass or low overflow metabolism). Case study 1 thus highlights NAViFluX’s ability to transform numerical flux distributions into pathway-level, mechanistically interpretable views of metabolism.

An important theme is that NAViFluX is not just a simple pathway viewer but also actively supports iterative model correction and analytic computation. In case study 2, visually identifying unrealistic bypasses (e.g. R15BPK) enabled adjustments of reaction constraints to recover experimentally supported essential genes (e.g. PRPPS) thereby modestly improving model performance when compared to KEIO reference knockout sets. Thus, by providing constraint refinement, NAViFluX enables GSMNs to more accurately capture real cellular metabolism. Similarly, visualizing merged subnetworks spanning glycolysis and PPP clarified the joint control of oxidative and non-oxidative PPP on *prpp* availability, suggesting RPI and TKT1 as rational targets for combinatorial perturbations. Thus, NAViFluX can be used as a sandbox where users can test ideas including alternative constraints, reaction additions / deletions or knockouts and immediately assess the consequence at the pathway level.

Case study 3 shows that NAViFluX is not only suitable for understanding physiological states but also for exploring engineered metabolic states. By adding CBB cycle reactions, constraining certain native reactions and simulating combinatorial uptake of pyruvate with glycerol, the platform was not only able to reproduce previously reported hemiautotrophic behavior but also maximize carbon fixation in *E. coli.* This demonstrates that NAViFluX can help design, interpret engineered flux modules and determine the trade-offs between growth, carbon catabolism and by-product secretion. Taken together with the first two case studies, NAViFluX provides a common environment for interrogating native physiology and experimenting with rational design for engineering systems.

As NAViFluX unifies such complex analyses within a single platform to provide biologically interpretable solutions, we envision it to become a routine tool in experimental and computational labs alike, particularly for exploratory hypotheses generation and for communication of model simulation results in logical, visual and flat-file output formats to non-computational collaborators.

### Limitations

NAViFluX facilitates merging of subsystems with flexibility of layouts. However, visualization performance may get affected for very large, merged networks. The platform does not yet support the automated reconstruction of networks from genomic information. Future work could therefore include visualization of community or microbiome models, integration with upcoming, more accurate representations of metabolism [36] and integration with automatic model reconstruction tools that can generate reconstructions directly from genomes.

## ACKNOWLEDGEMENTS

M.B.K acknowledges the support provided by Indian Council of Medical Research, Government of India for his Junior Research Fellowship. A.S., M.B.K and P.S.H. additionally thank IIT Hyderabad for the infrastructure support.

## AUTHOR CONTRIBUTIONS

Manjunatha Beduru Krishnamurthy: Formal analysis, Methodology, Validation, Writing. P S Harish: Formal analysis, Methodology. Abhishek Subramanian: Conceptualization, Formal analysis, Visualization, Writing

## DATA AVAILABILITY

All source code and supplementary files associated with the case studies are publicly available via Zenodo at https://zenodo.org/records/18137089. In addition, the complete source code, detailed documentation, test datasets, and step-by-step installation instructions are accessible through the project’s GitHub repository at https://github.com/bnsb-lab-iith/NAViFluX.

## SUPPLEMENTARY FILES

**Supplementary Figure S1.** Comparison of experimentally identified and NAViFluX-predicted flux-active metabolites. Venn diagram showing similarity between the experimentally detected metabolites and flux-active metabolites predicted by NAViFluX across the 11 carbon sources. The outer circle represents flux-active metabolites predicted by FVA implemented in NAViFluX. The inner circle represents the set of experimentally detected metabolites. Interestingly, these metabolites form a subset of the computationally predicted flux-active metabolites showing complete overlap suggesting strength in model predictions.

**Supplementary Table S1:** This supplementary file contains the Flux Variability Analysis (FVA) profiles for the *Escherichia coli* iJO1366 GSMN simulated under different carbon source conditions (Case study 1).

**Supplementary Table S2.** Over-representation analysis (ORA) results on the reactions obtained from the NAViFluX FVA functionality simulated for each Carbon source (Case Study 1)

**Supplementary Table S3.** Cycle-free flux balance analysis (cFBA) profiles for global, wild-type and specific reaction knockout simulations in *Escherichia coli* iJO1366 (Case study 2).

**Supplementary Table S4.** Flux profiles of engineered hemiautotrophic *E. coli* GSMN simulated under various carbon source conditions.

