## Supplementary Figure S1 for "NAViFluX: a visualization-centric platform for interactive analysis, refinement and design of genome-scale metabolic networks"

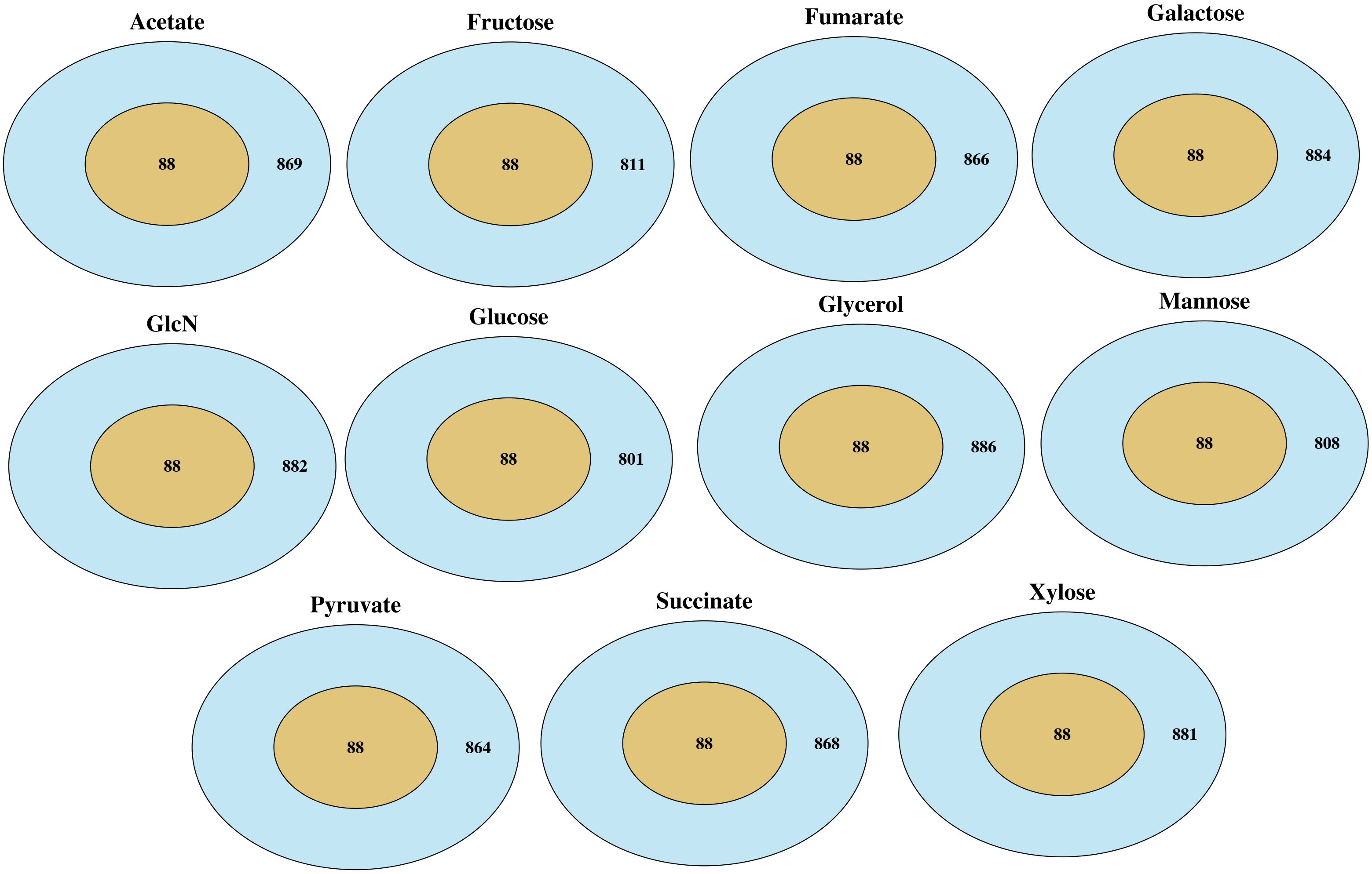


**Supplementary Figure S1.** Comparison of experimentally identified and NAViFluX-predicted flux-active metabolites
